# A multi-scale vascular atlas of blood vessels within the urinary bladder of male and female mice

**DOI:** 10.1101/2025.05.18.654773

**Authors:** LD Bowden, M Daglas, PB Osborne, JR Keast

**Affiliations:** Department of Anatomy and Physiology, University of Melbourne, Parkville VIC 3010 PDE

## Abstract

The vascular supply of the urinary bladder is embedded within a highly dynamic environment that includes alternating cycles of regional compression or stretching during bladder filling, sustained continence and voiding. These place unique demands on the vasculature to maintain tissue perfusion, fluid homeostasis and immune surveillance. Understanding this vascular regulation is also highly relevant to defining mechanisms of organ reperfusion following pelvic surgery, pelvic venous insufficiency, and the impacts of diabetes and ischemia on urinary function. There is limited anatomical knowledge on the organization of this vascular network, so we aimed to determine if there are stereotypical features associated with the mouse urinary bladder. We applied advanced microscopy and anatomical visualization methods to samples of the entire bladder viewed as a whole mount, including intravital tomato lectin labeling of the arterial vasculature, multi-channel immunofluorescence, tissue clearing, light-sheet and confocal microscopy. We developed a comprehensive multi-scale 3D anatomical map of the stereotypical arterial and venous networks associated with the mouse urinary bladder in both sexes, showing that the primary features of this network are established by the early postnatal period, prior to maturation of voiding and continence reflexes. These outcomes provide the foundation for probing mechanisms that underpin physiological and pathophysiological changes in the urinary bladder vascular network and a resource to guide more refined experimental perturbation, analysis and interpretation of vascular function/dysfunction in mouse models. This new knowledge on the structure of the urinary bladder vascular network will also benefit tissue engineering efforts seeking to restore or replace this organ.

## Introduction

Organs of the lower urinary tract (LUT)—the urinary bladder and urethra, with the associated striated muscle urethral rhabdosphincter—are tightly regulated by the nervous system to initiate voiding (micturition) and storage (continence) at specific times (1, 2). The blood vessel network of these organs is essential for appropriate tissue perfusion, fluid homeostasis, and immune surveillance but is embedded within a highly dynamic environment. This includes extreme alternating cycles of regional compression or stretching during bladder filling, sustained continence with increased sphincter tone, and voiding. This places specific demands on the LUT vasculature and its neural regulation (3, 4), which align with unique adaptive features of the LUT vasculature that contribute to both episodic and longer-term adjustments of vascular flow: including structural specializations (5–11), intrinsic non-neural and extrinsic neural mechanisms (4, 12, 13). However, to place these in the context of overall vascular regulation it is essential to understand the anatomical organization of this network.

To our knowledge, the source and potential stereotypical patterning of blood vessels within the urinary bladder have not been described. This is a significant limitation in the context of research determining how vascular dysfunction is involved in a diverse range of clinical situations. These include organ reperfusion following pelvic surgery, pelvic venous insufficiency and pelvic congestion syndrome (a chronic pain condition involving the pelvic venous system), bladder cancer, diabetes and ischemia, a likely contributor to many conditions, including overactive bladder (12, 14–22, 22, 23). Determining the structure of this vascular network will also benefit tissue engineering efforts seeking to restore or replace LUT organs (24–26).

Our aim was to define the structural organization of the blood vessel networks associated with the urinary bladder of adult mice, by identifying the primary vascular sources and their networks within specific regions and tissues. We also wished to determine the primary features of this network within the early postnatal period, when voiding and continence mechanisms have not yet fully matured (1).

The multi-scale nature of this goal, along with the three-dimensional (3D) and complex branching nature of the vascular network, limits the information that can be obtained from conventional histological sections. We addressed this technical challenge by applying tissue clearing methods (27) to intact whole mounts of the entire urinary bladder viewed with light sheet microscopy and sub-dissected, full thickness opened (flat mount) preparations viewed with confocal microscopy. Our study also incorporated a protocol for intravital labeling with tomato lectin (28) that enabled us to selectively view the arterial network, using immunofluorescence to visualize the entire blood vessel network.

Together these approaches have enabled us to build a comprehensive multiscale 3D understanding of arterial and venous networks and capillary beds across the urinary bladder in young adult mice (postnatal days 49-70) and pups (P7-10). This knowledge base has been developed for both male and female mice, due to the unique relationships between the LUT and adjacent reproductive organs (prostate gland, uterine cervix and vagina) and allowing identification of any sex differences within the vasculature. All images underpinning our study will be publicly available for further analysis (sparc.science).

## Materials and Methods

### Animals

The primary investigations of this study used 27 female and 18 male adult C57BL/6J mice (7-10 weeks, females 16-21 g, males 20-25 g). Mice were sourced from the Animal Resources Centre (Ozgene; Bentley, WA, Australia) and Australian BioResources (Moss Vale, NSW, Australia).

The estrus cycle of all adult females was recorded. For studies on pups, 16 female and 15 males were used (7-10 days, females 2.6-7.0 g, males 3.0-6.5g). Pups were acquired by purchasing time-mated dams or breeding in house.

All procedures were approved by the University of Melbourne Animal Ethics Committee in accordance with the Australian Code for the Care and Use of Animals for Scientific Purposes (National Health and Medical Research Council of Australia). Adult mice of the same sex were housed in groups of 2-4 under a 12-hour light-dark cycle with free access to food and water.

Pups were housed with their mother until the time of experimentation.

### Tissue collection

Mice were anesthetized (150 mg/kg ketamine and 15 mg/kg xylazine, i.p.) prior to intracardiac perfusion. The perfusion method differed for studies where the bladder was fixed intact (referred to as intact whole organs) and studies where the bladder was dissected separately from the urethra, cut open, pinned flat and fixed (opened flat mounts). Opened flat mounts enabled specific features to be viewed with confocal microscopy at a higher resolution than available using light sheet microscopy. In most animals of both groups, the first step following anesthesia and opening of the chest cavity was injection of conjugated *Lycopersicon esculentum* (Tomato) lectin (1 mg/ml) into the left ventricle of the heart using 31G needle over 30 seconds (100 µl in adults, 20-50 µl in pups) (28). The lectin conjugates used for this study are described in **Table 1**. After waiting a minute (or 40s in pups), intracardiac perfusion was performed as described below. This protocol labels the arterial and capillary but not venous networks. A vaginal smear was taken from each adult female mouse to assess the estrus cycle.

**Table 1.**
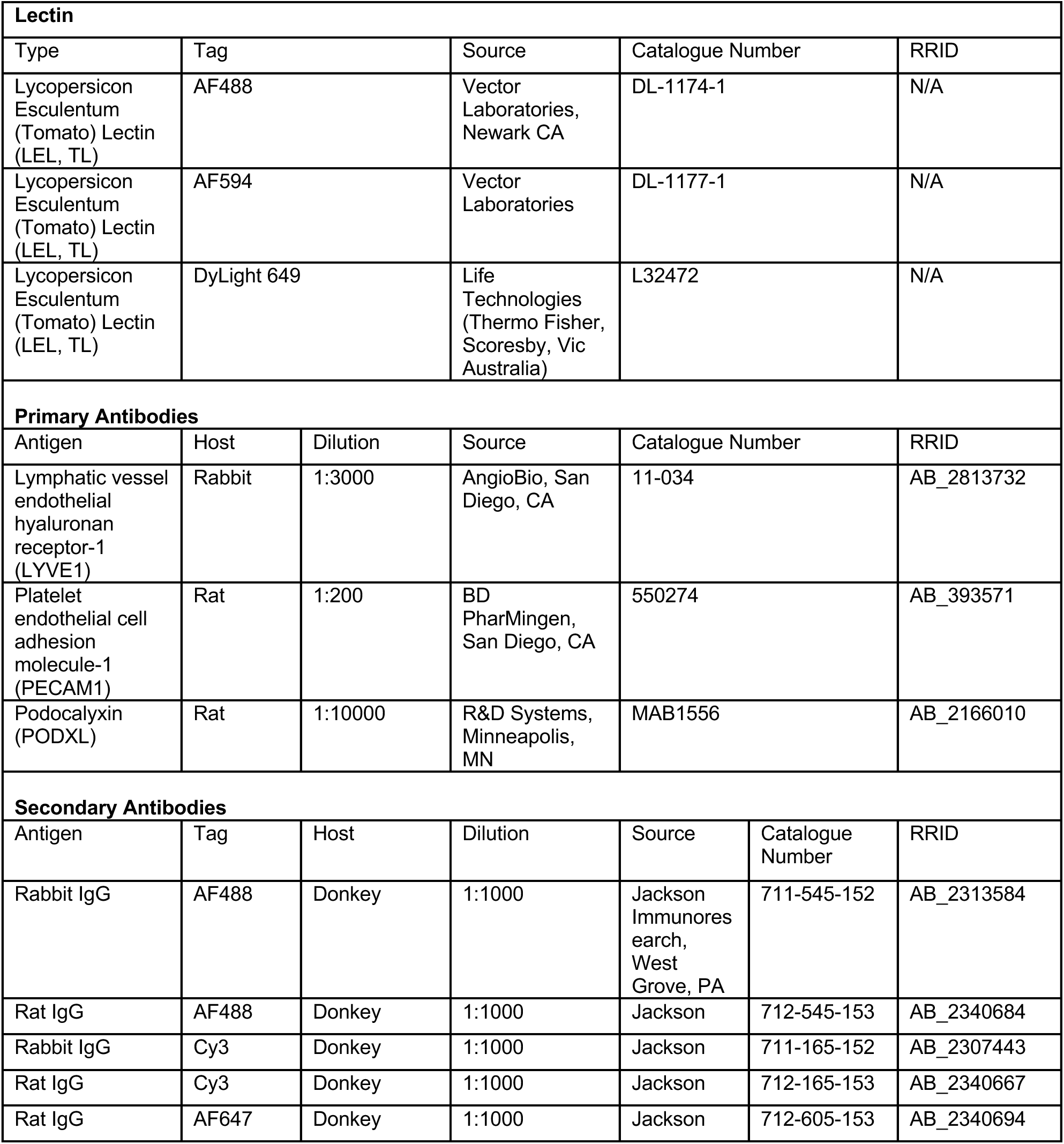
Antibodies and lectins.

### Preparation of intact whole organ samples

Anesthetized adult mice (18 female, 14 male) were perfused intra-cardially using 18 ml of pre-wash (1% sodium nitrite and 5000UI/ml heparin in 0.9% saline) followed by 90 ml of fixative (4% paraformaldehyde in 0.1 M phosphate buffer, pH 7.4) at the rate of 18 ml/min. Pups (16 female, 15 male) were perfused with 7.5 ml pre-wash, followed by 15 ml fixative at a rate of 3 ml/min.

The entire LUT was removed and postfixed overnight in 4% PFA, followed by three 30-minute washes in 0.1M phosphate buffered saline (PBS, pH 7.2). Tissue was stored in PBS with 0.1% sodium azide (PBS-azide) at 4°C. In six adult samples (3 female, 3 male) the entire pelvis was left intact. In this study, only the urinary bladder and its vasculature were analyzed. The vasculature of the attached urethra and sphincter was highly complex, required additional approaches and will be described in a separate publication.

### Preparation of opened flat mount samples

Opened flat mount samples of urinary bladder were prepared from adult mice (9 female, 4 male). Instead of fixative, intracardiac perfusion was performed only with prewash 90 ml, at the rate of 18 ml/min, after which the bladder with short sections of attached proximal urethra and ureters was dissected. The tissue was sub-dissected further while submerged in a prewash-filled petri dish with a silicon polymer base. To produce the flat mount, the bladder was first cut open along either the ventral or dorsal midline, depending on the purpose of the preparation (i.e., which group of midline structures were to remain intact). The bladder was then pinned out flat in the petri dish using pins (0.0125 mm diameter) and fixed overnight (4% PFA), washed three times for 30 minutes in 0.1M PBS and stored in PBS-azide at 4°C.

### Immunolabelling and tissue clearing of intact whole organ samples

The tissue clearing protocol was based on iDISCO (27). Samples were washed in 1x Dulbecco’s PBS (DPBS; Sigma-Aldrich, NSW, Australia) six times for 15 minutes, then dehydrated in increasing concentrations of methanol for 1.5 hours each (50% and 80% in DPBS, 100%) and incubated overnight in 6% hydrogen peroxide in methanol at 4°C. Following this, samples were rehydrated in decreasing concentrations of methanol for 1.5 hours each (100%, 100%, 80%, 50%) then incubated in DPBS for 1.5 hours. The samples were then transferred into blocking solution for 3 (pups) or 4 days (adults) at room temperature (DPBSG-T; DPBS with 0.2% gelatin, 0.5% Triton-X and 0.01% thimerosal). They were then incubated in DPBSG-T containing primary antibodies for 10-14 days at 37°C, then washed six times for 15 minutes (DPBS-T; DPBS with 0.5% Triton X-100). Samples were then incubated in secondary antibodies in DPBSG-T for 2 days at 37°C, then washed six times for 1 hour in DPBS-T. Some of the smaller samples from pups were embedded in agarose prior to the next dehydration step to facilitate specific orientation later within the microscope chamber. All samples were then dehydrated in increasing methanol concentrations in DPBS for 1 hour (20%, 40%, 60%, 80%, 100%, 100%). This stage was extended for the complete pelvis preparations, where each step was 2 hours, except for the 80% step (overnight incubation). Samples were then incubated overnight in 66% dichloromethane (DCM) and 33% methanol. Finally, samples were incubated in 100% DCM for 30 minutes, repeating this step until they had sunk in fresh DCM, then optically cleared in dibenzyl ether (DBE) for 2 hours and stored at room temperature in fresh DBE. All stages were performed at room temperature unless otherwise stated. Samples were constantly rotated throughout all incubations and washes.

Some samples included bone and were decalcified prior to iDISCO tissue clearing. This followed a previously published method (29). Samples dissected from pups with pubic bone attached and sectioned were incubated in 15 ml 0.5M ethylenediaminetetraacetic acid (EDTA, pH 7.4) for two days, changing to fresh EDTA solution on the second day. Samples of the entire pelvis dissected from adult mice were decalcified incubating in 50 ml 0.5M EDTA for 7 days, changing the solution on the morning of the fourth day.

Details of primary and secondary antibodies are provided in **Table 1**. To view the total population of blood vessels, antibodies to platelet endothelial cell adhesion molecule-1 (PECAM1; also known as CD31) and podocalyxin (PODXL) were combined and visualized using the same conjugated secondary antibody. This method enhances the quality of vessel labelling (30).

### Immunolabelling of opened flat mount samples

Opened flat mount preparations of urinary bladder were washed in 0.1M phosphate buffer (PB, pH 7.4) three times for 30 minutes, then twice for 30 minutes in 50% ethanol/H_2_O and incubated in blocking solution (PB containing 10% non-immune horse serum (NHS)) for 30 minutes. The tissues were then incubated for 3 days in primary antibody solution (0.1% PBS-azide, 0.5% Triton X-100, 10% NHS). Flat mounts were washed three times for 30 minutes in PBS then incubated in secondary antibodies (0.1% PBS-azide, 0.5% Triton X-100,10% NHS) for 3 days. They were then washed in PBS three times for 30 minutes, then for 1 hour in mounting medium (buffered glycerol, pH 8.6). Finally, the flat mounts were mounted on a glass slide and cover-slipped with buffered glycerol. Tissues were mounted with serosa or lamina propria side up, depending on the specific imaging required. Slides were sealed with nail polish and stored at 4°C. All steps were completed at room temperature on rotation. Details of primary and secondary antibodies are provided in **Table 1**.

### 3D imaging of intact whole organs (light sheet microscopy)

Cleared organs were transferred to ethyl cinnamate and imaged using light sheet fluorescence microscopy (Ultramicroscope II or Ultramicroscope Blaze, Miltenyi Biotec, Germany). Samples imaged on the Ultramicroscope II were imaged using a zoomable 2x lens (MVPLAPO, Olympus, Japan) with the numerical aperture of the light sheet set to 0.156. Samples imaged using the Ultramicroscope Blaze used either 1.1x objective (1.1x/0.1 MI PLAN), 4x objective (4x/0.35 MI PLAN) or 12x (12x/0.53 MI PLAN) with alternate zooms of 0.6x, 1x, 1.66x and 2.5x. The numerical aperture of the light sheet was set to 0.163. For both microscopes, single-sided and dual-sided 3-sheet illumination was used with lasers 488, 561 and 639 paired with emission filters 525/50, 620/60 and 680/30, respectively. The exposure times were 50 – 200 ms. The z-steps were 2, 5 or 10 µm, depending on the purpose of the image (i.e., segmentation, higher quality overview and basic overview images, respectively). Any tiled images were taken with 10% overlap. Image stacks were converted to Imaris file types (Imaris File Converter, Bitplane) and if the image was a mosaic, stitched using Imaris Stitcher (Bitplane).

### Imaging of opened flat mounts (wide field and confocal microscopy)

Overview images of opened flat mount preparations were taken using widefield (5x Plan-Apochromat, Zeiss AxioImager M2, Carl Zeiss Microscopy). Confocal microscopy was used to image targeted structures (10x Plan-Apochromat, Zeiss LSM800 Airyscan, Carl Zeiss Microscopy; 10x, 20x and 40x Plan-Apochromat, Zeiss LSM900, Carl Zeiss Microscopy; 10x and 20x Plan-Apochromat, Zeiss LSM900 Airyscan2, Carl Zeiss Microscopy). Mosaic images were stitched using Zen Blue (Zeiss). Some of these images were deconvolved to visualize specific structures more clearly (deconvolution package, Zen Blue, Zeiss).

### Visualization and qualitative analysis of vascular structures

The light sheet images of the entire bladder were viewed using Imaris (Bitplane). Overview scans were first performed to assess the quality of the preparation (organ and vascular completeness to indicate dissection quality; signal and clarity of lectin and immunolabeling to indicate processing quality). Typically, autofluorescence of the shorter wavelength channel (488) allowed for visualization of the organs. The smaller z-step images were used to visualize the structures and complete segmentation of key objects of interest. Arteries were traced in Imaris using the *filament tracer* function to highlight structures visible in the lectin channel. When specific regions of the vasculature showed inconsistent lectin labelling, the endothelial marker channel was used. In some cases, the *surface* function was used to visualise the serosal surface of organs or remove artifacts (such as bubbles or urine crystals) to aid in the visualization of key structures. When using the *surface* function to outline an organ, it was manually drawn every 50 z-steps in the 2D plane using the autofluorescence signal (488 channel). In addition to the light sheet images, some widefield and confocal images underwent segmentation of specific objects using the *surface* function.

For the cleared pelvis samples an alternate approach to segmentation was used. Visualization and segmentation occurred using virtual reality software syGlass (IstoVisio, Inc., Morgantown WV) coupled with an Oculus Quest2 (Meta) headset. Using the ROI tool, each vessel was segmented and then exported as tiff files. A custom python code was then used to extract the segmentation completed from the raw data (29). The output provides the underlying fluorescence of each mask created as independent files. These tiff stacks were then converted back into Imaris files and imported as additional images in the original Imaris file.

### Figure production

All mapping studies were performed on multiple replicates from each sex but have been illustrated in both sexes only where there was an identified difference. Widefield and confocal images were typically adjusted (levels) to best reflect what was seen under the microscope. Images were taken using the snapshot function in Imaris or exported as an image using Zen Blue. To enhance visibility of vascular structures, some images have been provided as monochrome with inverted grey scale. Minor adjustments in levels (Photoshop, Adobe Creative Suite) have been made in some images to improve visibility of delicate structures. Videos of 3D light microscopy images were created using the video function in Imaris and annotated using Premier Pro (Adobe Creative Suite). Schematics were created in Illustrator and figures were constructed in InDesign (Adobe Creative Suite).

## Data availability

Original image files and 3D visualizations (segmentations) will be available for viewing or download on sparc.science. Videos are provided as Supplementary files available in Figshare.

## RESULTS

### Visualization of the blood vascular system

The intact, whole lower urinary tract (LUT), including the associated urethral rhabdosphincter, was cleared (Fig. 1A) prior to light sheet microscopy. To provide a macroscopic context for more detailed analysis of intramural network components, we visualized the extramural sources of the urinary bladder arterial network in three female and three male adult mice. In these dissections, many surrounding tissues (including pelvic bones) were retained to ensure the relevant major vessels were not damaged. In the larger samples including pelvic bones, the pelvic striated muscles could be readily identified by their autofluorescence (Fig. 1B).

**Figure 1.**
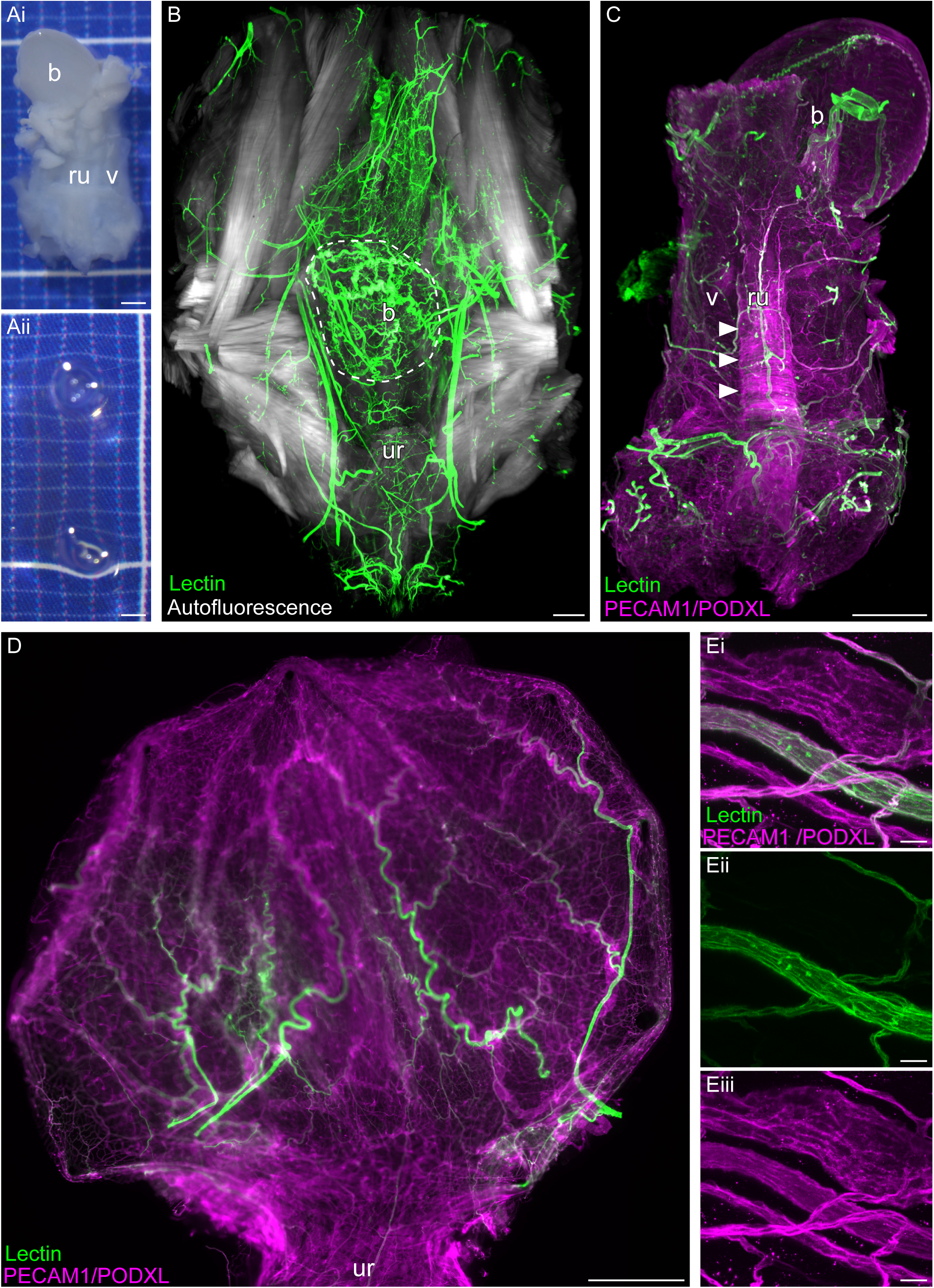
Visualization of vascular networks in different types of lower urinary tract (LUT) preparations of the adult mouse. To visualize the vasculature of the entire, intact bladder, the bladder was dissected with surrounding tissues still attached, then cleared and immunolabelled (**A-C**). For higher resolution analysis of the vasculature, opened flat mount preparations were used (**D, E**). All samples shown were from female mice. Images **A-C** are oriented from the ventral view with bladder at the top of the image. Images **B-E** show samples containing fluorophore-conjugated tomato lectin. When administered intra-cardially, this lectin labelled arterial and capillary but not venous networks. b, bladder; ru, rhabdosphincter and urethra; ur, urethra; v, ventral vaginal wall. **A**, Entire, fixed LUT before and after tissue clearing, viewed with a dissecting microscope. Grid units = 1 mm. **B**, Light sheet image of the LUT in a sample where many of the surrounding pelvic structures were retained. This enables visualization of the entire extramural supply to the bladder and surrounding pelvic structures. Autofluorescence demonstrates the structure of pelvic striated muscles. In this sample the bladder is folded over towards the viewer (boundary indicated by the dotted line), slightly obscuring the adjacent urethra. **C**, Light sheet image of a cleared, intact LUT where most extramural vessels and tissues have been removed, making it easier to visualize the vasculature specifically associated with and located within the LUT organs. This demonstrates the arterial system (lectin) and total blood vascular system using a combination of antibodies against platelet endothelial cell adhesion molecule-1 (PECAM1) and podocalyxin (PODXL). Autofluorescence associated with the rhabdosphincter is also visible in the PECAM1/PODXL channel; one edge of the sphincter is indicated with arrowheads. **D-E:** Flat mount of the urinary bladder opened along the dorsal midline prior to fixation, viewed with wide field (**D**) or confocal microscopy (**E**). Panel **D** is oriented with the bladder apex at the top of the image, lateral margins are the dorsal midline; midline of the image is the ventral midline of the bladder. The venous network can be identified by comparing tomato lectin labelling (arterial and capillary) with PECAM1/PODXL immunolabeling (all blood vessels), where venous structures are PECAM1/PODXL-positive and lectin-negative. Scale bars: A-D, 1000 µm; E, 20 µm.

Intracardiac injection of fluorescently tagged tomato lectin into anesthetized adult mice prior to perfusion with fixative enabled direct visualization of the arterial network and capillaries (Fig. 1 **B-E**). When the entire blood vascular network was visualized by immunolabeling for endothelial proteins (PECAM1 and PODXL), the venous network could be identified by the absence of lectin (Fig. 1E).

### Arterial network of the adult urinary bladder

In both male and female mice, we identified four distinct arterial vessels that supply the adult urinary bladder. We named these vessels by their stereotypical location and trajectory: superficial ventrolateral artery (SVLA), deep dorsal artery (DDA), deep ventral artery (DVA) and dorsal midline artery (DMA). These will each be described in detail below and illustrated in Figures 2-4**, Supplementary Videos S1-S4** (S1 https://doi.org/10.26188/28844174.v1 S2 https://doi.org/10.26188/28844177.v1 S3 https://doi.org/10.26188/29083577.v1 ; S4 https://doi.org/10.26188/29062364.v1), with the number of replicates and anatomical variations provided in **Table 2** and summary schematics in Figure 6. These schematics include the primary anatomical variations described below.

**Figure 2.**
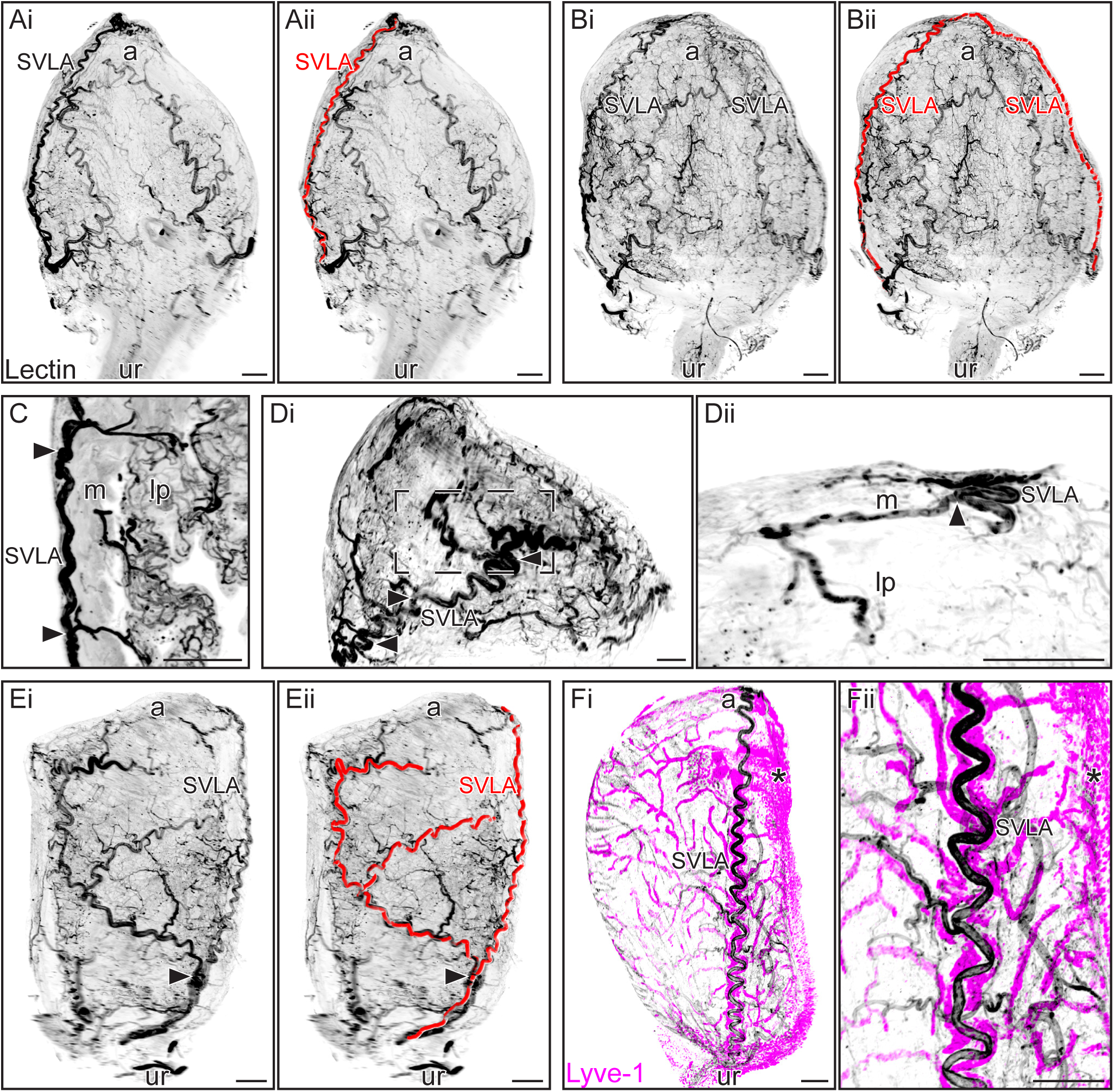
The superficial ventrolateral artery (SVLA) of the adult mouse urinary bladder. Light sheet microscopy of cleared intact whole bladders dissected from female mice. Intracardiac injection of fluorophore-conjugated tomato lectin preceded fixation. Specific regions and tissue layers are indicated: apex (a), urethra opening (ur), lamina propria (lp) and muscularis (m). Because the samples have been cleared, vessels throughout the wall of the bladder are visible. All segmentations were performed with Imaris. Monochrome images have been inverted to enable better visualization of vascular structures. **A**, Ventral view of the bladder, showing the SVLA on the left side of the image (right side of the bladder). Original image (**Ai**) shows the SVLA and many other arterial vessels; the SVLA has been segmented in **Aii**. **B**, Ventral view of the bladder from a mouse that had two SVLAs, one on each ventrolateral side (**Bi**, original; **Bii**, SVLA segmented). **C**, Optical slice (150 µm) of a region of SVLA, arrows indicating branches supplying the capillary bed in the muscularis (m) and lamina propria (lp). **D**, Superior view of the bladder showing the region near the apex of the bladder, oriented with the ventral region at the bottom of the image. **Di**, The SVLA (arrows) is highly tortuous near the apex. Inset is magnified in **Dii**, where an optical slice (300 µm) of the SVLA has been provided to show its branches entering the muscularis and lamina propria. **E**, Lateral view of the bladder where the SVLA had a laterally projecting branch; branch point indicated by an arrow (**Ei**, original; **Eii**, SVLA segmented). Image oriented with ventral on the right, showing bifurcation of the SVLA on the dorsal side. **F**, Lateral view of the bladder showing the SVLA surrounded by lymphatic vessels identified by Lyve-1 immunoreactivity and at two levels of magnification). Lyve-1 positive macrophages indicated by asterisk. Scale bars: 300 µm.

**Table 2:**
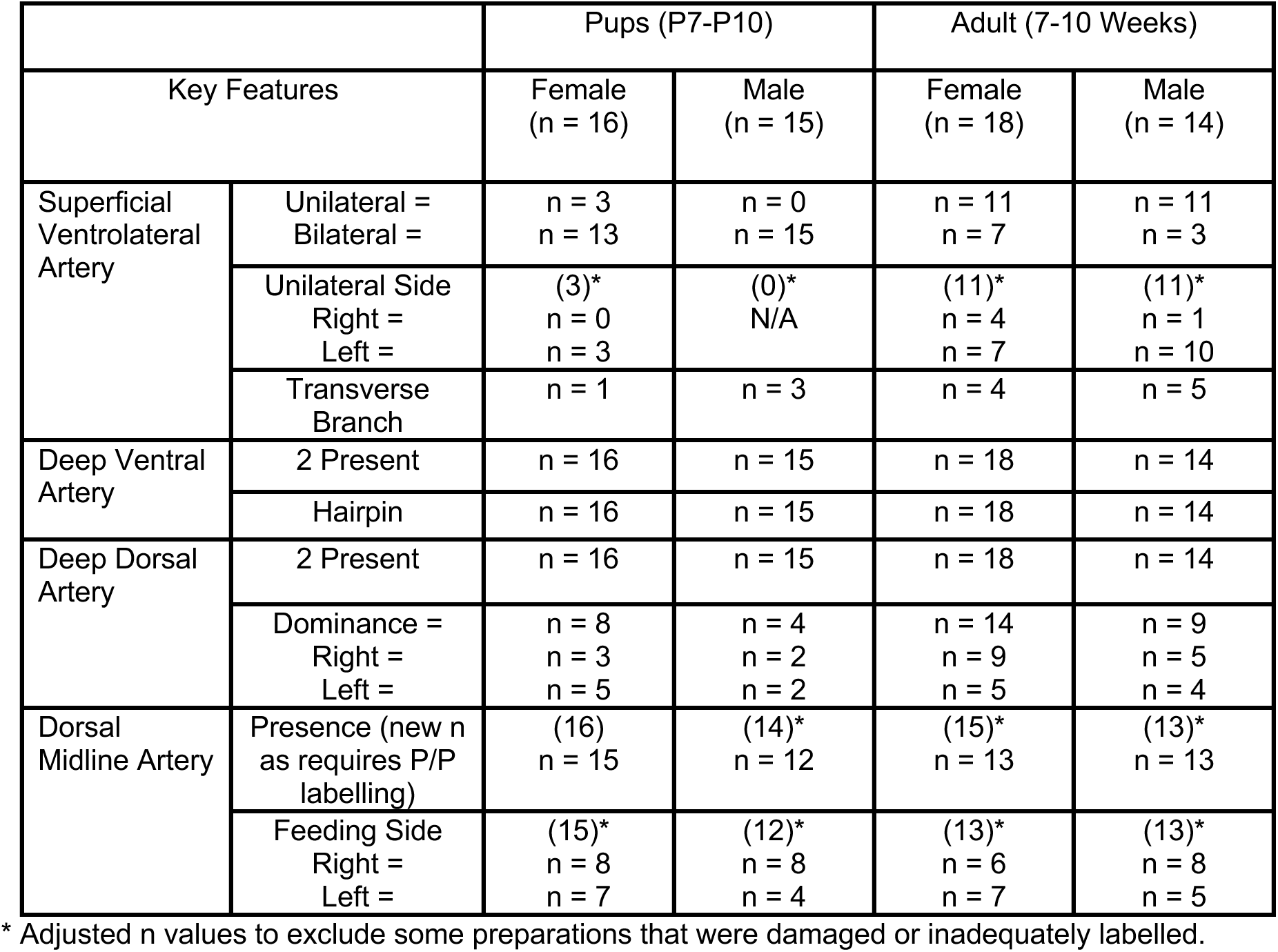
Primary features and anatomical variations of bladder arterial vessels.

|  |  | Pups (P7-P10) |  | Adult (7-10 Weeks) |  |
| --- | --- | --- | --- | --- | --- |
| Key Features |  | Female<br>(n = 16) | Male<br>(n = 15) | Female<br>(n = 18) | Male<br>(n = 14) |
| Superficial Ventrolateral Artery | Unilateral =<br>Bilateral = | n = 3<br>n = 13 | n = 0<br>n = 15 | n = 11<br>n = 7 | n = 11<br>n = 3 |
|  | Unilateral Side<br>Right =<br>Left = | (3)*<br>n = 0<br>n = 3 | (0)*<br>N/A | (11)*<br>n = 4<br>n = 7 | (11)*<br>n = 1<br>n = 10 |
|  | Transverse Branch | n = 1 | n = 3 | n = 4 | n = 5 |
| Deep Ventral Artery | 2 Present | n = 16 | n = 15 | n = 18 | n = 14 |
|  | Hairpin | n = 16 | n = 15 | n = 18 | n = 14 |
| Deep Dorsal Artery | 2 Present | n = 16 | n = 15 | n = 18 | n = 14 |
|  | Dominance =<br>Right =<br>Left = | n = 8<br>n = 3<br>n = 5 | n = 4<br>n = 2<br>n = 2 | n = 14<br>n = 9<br>n = 5 | n = 9<br>n = 5<br>n = 4 |
| Dorsal Midline Artery | Presence (new n as requires P/P labelling) | (16)<br>n = 15 | (14)*<br>n = 12 | (15)*<br>n = 13 | (13)*<br>n = 13 |
|  | Feeding Side<br>Right =<br>Left = | (15)*<br>n = 8<br>n = 7 | (12)*<br>n = 8<br>n = 4 | (13)*<br>n = 6<br>n = 7 | (13)*<br>n = 8<br>n = 5 |

\* Adjusted n values to exclude some preparations that were damaged or inadequately labelled.

The SVLA (Fig. 2A**-C**) joined the bladder near the bladder neck, then projected ventrolaterally on the serosal surface to the apex. The SVLA was either a single structure (11/18 females, 11/14 males) or bilateral (7/18 females, 3/14 males). Examples are shown in Figure 2A (single) and Figure 2B (bilateral). Along this trajectory, branches from the SVLA formed capillary beds in the underlying muscularis and lamina propria (Fig. 2C). These branches were sparse until the SVLA reached the region close to the apex, where the SVLA became more tortuous and branched more profusely to form extensive capillary beds in both tissue layers (Fig. 2D).

We further examined the anatomical variations in the SVLA and found that when unilateral, it was most commonly present on the left side of the bladder (7/11 females, 10/11 males). We also found that in a minority of mice (4/18 females, 5/14 males), the SVLA had a large branch projecting towards the lateral region of the bladder. This branch occurred approximately midway along the length of the SVLA and bifurcated on the dorsal side of the bladder (Fig. 2E). This was seen in mice that had either one or two SVLAs, so overall was more commonly present on the left side. This branch typically travelled between the muscularis and lamina propria to supply capillaries to both the tissue layers.

In specimens immunolabelled for PECAM1 and PODXL to visualize the entire blood vascular system, weakly labelled vessels were identified on each side of the SVLA. Further investigation using Lyve-1 immunofluorescence identified that these comprise a pair of lymphatic vessels travelling on each side of the SVLA, following their entire length until reaching the apex of the bladder (Fig. 2F). When two SVLAs were present, each was closely accompanied by lymphatic vessels. These structures were investigated further when assessing the venous network (see below and Fig. 5). A thorough investigation of the lymphatic vasculature will be the focus of a separate study.

The deep ventral arteries (DVA, Fig. 3A) arose from their respective vesical arteries close to the bladder neck then immediately penetrated the bladder serosa and passed through the muscularis. Each DVA then projected along the boundary of the muscularis and lamina propria of the ventral bladder wall towards the apex, where it joined with the DVA from the other side to form a hairpin structure (Fig. 3Ai**-iii**). Along this interlaminar trajectory, perpendicular branches at regular intervals provided capillary beds of the muscularis and lamina propria (Fig. 3Aiii).

**Fig. 3.**
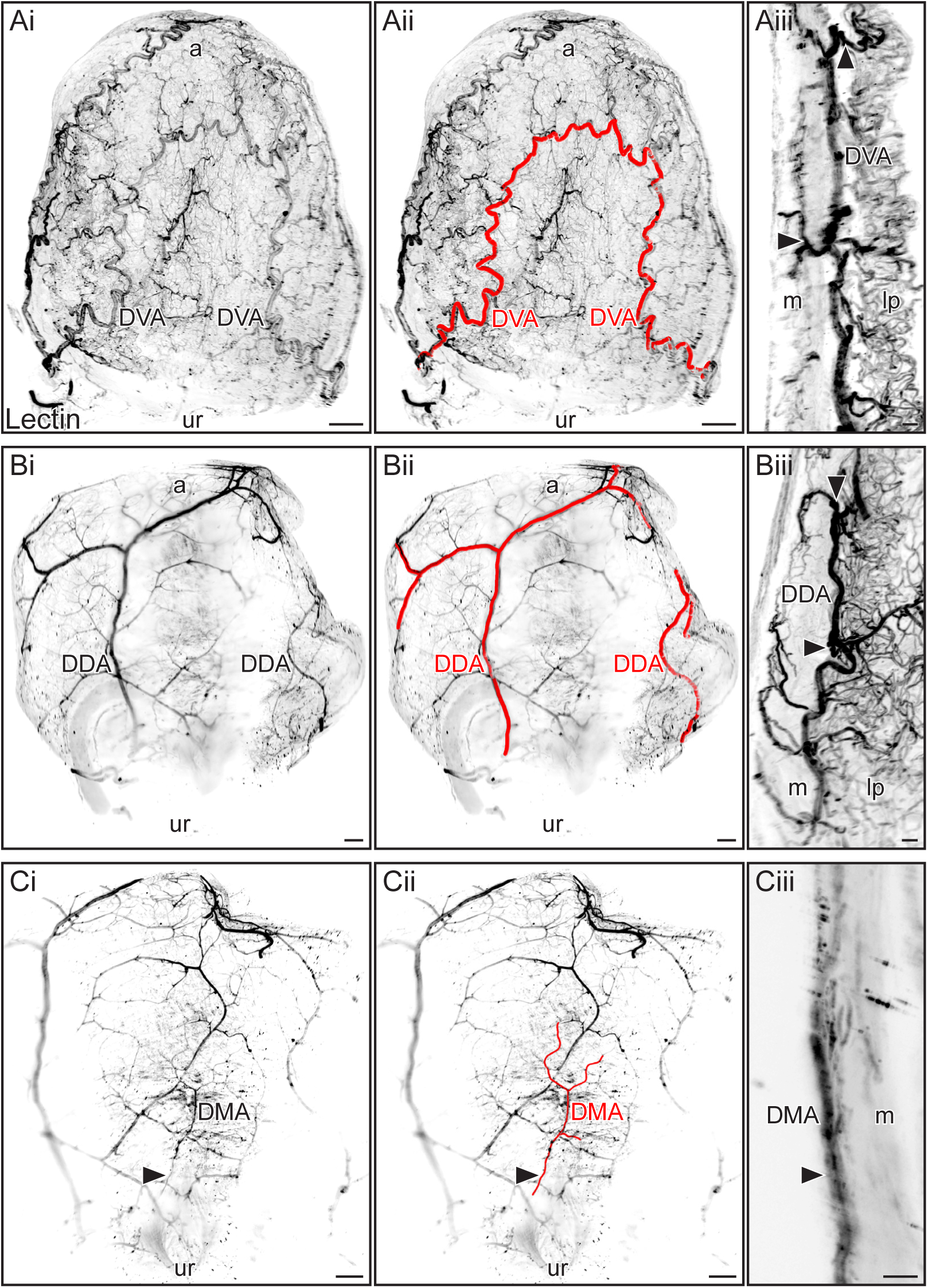
The deep ventral artery (DVA), deep dorsal artery (DDA) and dorsal midline artery (DMA) of the adult mouse urinary bladder. Light sheet microscopy of cleared whole bladders dissected from female (**A, C**) and male (**B**) mice. Intracardiac injection of fluorophore-conjugated tomato lectin preceded fixation and labelled arterial structures. Specific regions and tissue layers are indicated: apex (a), urethra opening (ur), lamina propria (lp) and muscularis (m). All segmentations were performed with Imaris. Monochrome images have been inverted to enable better visualization of vascular structures. **A**, Ventral view of the bladder showing pair of DVAs joining near the bladder apex to form a hairpin structure (**Ai**, original image; **Aii** DVAs segmented). **Aiii**, an optical slice (350 µm) showing branches of the DVAs supplying the muscularis and lamina propria capillary beds (examples indicated by arrows). **B**, Dorsal view of the bladder showing a pair of DDAs, with the left DDA supplying a large area on the dorsal side of the bladder (**Bi**, original image; **Bii** DDAs segmented). **Biii**, an optical slice (350 µm) showing branches of the DDAs supplying the muscularis and lamina propria capillary beds (examples indicated by arrows). **C**, Dorsal view of the bladder showing the DMA projecting along the dorsal midline of the bladder (**Ci**, original image; **Cii**, DMA segmented). **Ciii**, an optical slice (200 µm) showing the DMA on the serosal surface of the bladder (arrows). Scale bars: i, ii 300 µm; iii 50 µm.

Similarly to the DVA, upon reaching the bladder the deep dorsal arteries (DDA, Fig. 3B) immediately passed through the serosa and muscularis, then projected along the boundary of the muscularis and lamina propria of the dorsal bladder wall towards the apex. Similarly to the DDV, projecting along the interlaminar boundary to the apex; at regular intervals perpendicular branches provided capillary beds of the muscularis and lamina propria (Fig. 3Biii). However, in contrast to the DVAs that joined in a hairpin structure, each DDA remained separate and appeared to supply a distinct region of dorsal bladder. The area supplied by the DDA was asymmetric in some animals (manual inspection, no quantitation of area). In adult female mice, the right DDA more commonly supplied a larger area of the dorsal side of the bladder (9/18), than the left DDA (5/18), with 4 bladders having symmetrical supply from the DDAs. The same analysis in adult male mice showed that in 5/14 mice a larger area was supplied by the right DDA, 4/14 had a larger supply from the left DDA and in 5 mice the supply was symmetrical.

A fourth arterial supply, the dorsal midline artery (DMA) was identified in the majority but not all animals (Fig. 3C**, Supplementary Video S2** https://doi.org/10.26188/28844177.v1). This artery was difficult to observe due to its much smaller diameter and midline dorsal location where closely apposed organs made it more challenging to image deeper tissues. Where the lectin signals and imaging were of sufficient quality to assess this specific region, the DMA was identified in 13/15 females and 13/13 males. The DMA arose from one of the urogenital arteries, a vesical artery or an artery supplying the vagina in females and the prostate in males, originating on either side, with no clear preference (**Table 2**). In some animals the DMA showed a clear bifurcation (Fig. 3Ci**, ii**). The DMA entered the bladder near the bladder neck, where it projected for a short distance along the serosa, incompletely penetrating the muscularis where it provided some capillaries (Fig. 3Ciii), then stopping around midway between the bladder neck and apex.

Analysis of the entire cleared pelvis preparations in combination with lectin labelling made it possible to determine the extramural origin of the urinary bladder vascular supply. The observations described below apply to both sexes. The SVLA, DVA and DDA arose via one or two vesical arteries (VA). These VAs branched from the urogenital artery (Fig. 4) (**Supplementary Videos S3** https://doi.org/10.26188/29083577.v1 **S4** https://doi.org/10.26188/29062364.v1 ), which originated from the ventral (anterior) division of the internal iliac artery. In some mice, the number of VAs differed on each side, as shown in Fig. 4A, where one VA supplied the left and two VAs supplied the right side. In contrast, in the sample shown in **Fig 4B**, there is only one VA on each side. It was more difficult to fully trace the origin of the DMA due to the density of surrounding tissues but was found to have a more variable location of its origin. Varying between animals, the DMA originated from vesical artery, reproductive branch of the urogenital artery or the urogenital artery itself.

**Figure 4.**
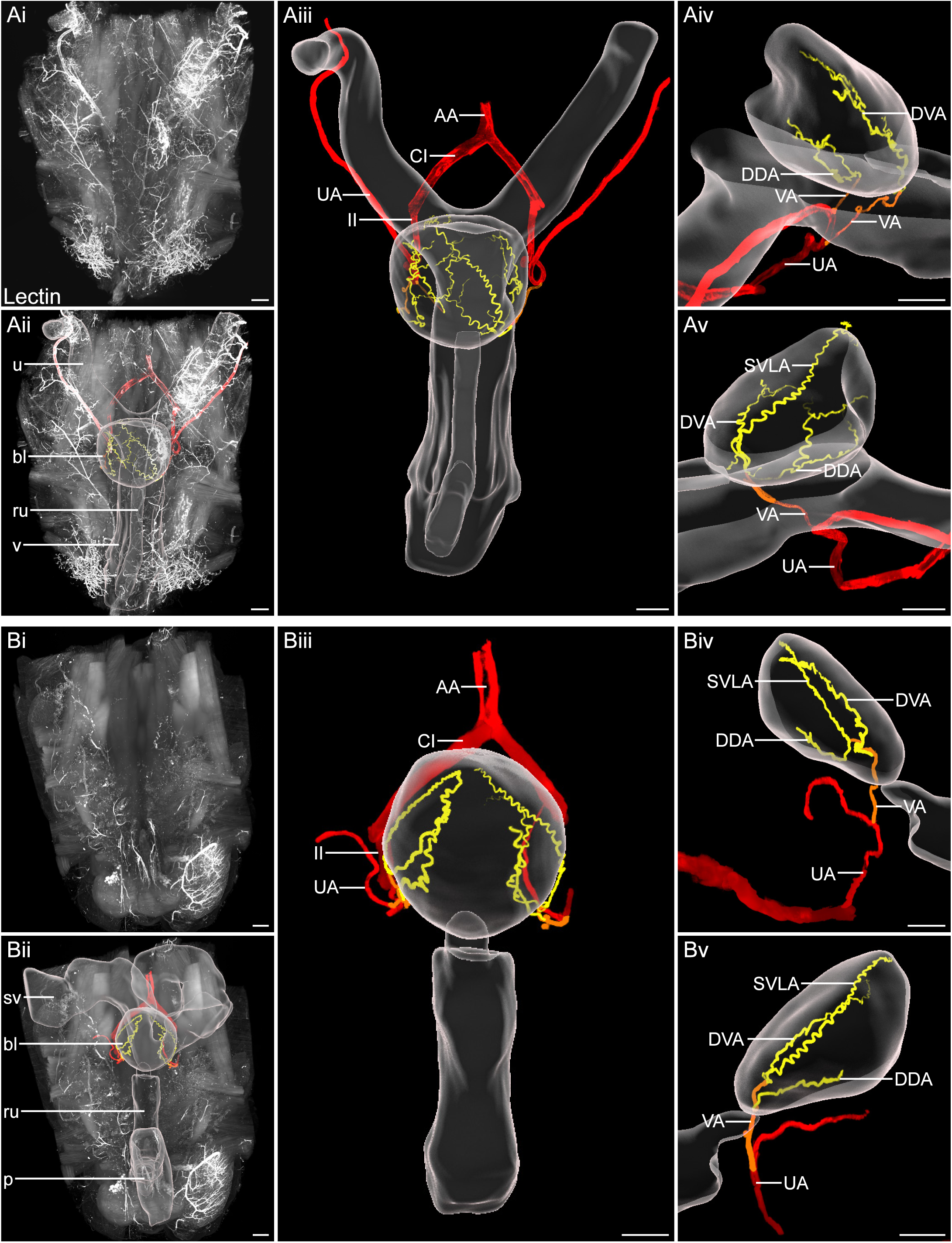
Location of macroscopic and extramural components of the adult mouse LUT vasculature. Light sheet microscopy of entire cleared pelvis preparations of female (**A**) and male (**B**) adult mice. Intracardiac injection of fluorophore-conjugated tomato lectin preceded fixation and labelled the arterial system. Organs surrounding the bladder are present within the sample. Segmentations of vessels and organs are completed using Syglass and Imaris respectively. Vessels in red are the vessels of the external supply (AA, abdominal aorta; CI, common iliac; II, internal iliac; UA, urogenital artery). The vesicular arteries (VA) are orange, and the vessels of the bladder are yellow (DDA, deep dorsal artery; DVA, deep ventral artery; SVLA, superficial ventrolateral artery). The dorsal midline artery is not shown as it was difficult to visualize and trace in these larger samples. Ventral view (**Ai-Aiii, Bi-Biii**) and lateral view (**Aiv, Av, Biv, Bv**). bl, bladder; p, penis; ru, rhabdosphincter and urethra; sv, seminal vesicles; u, uterus; v, vagina **A**, Female pelvis labelled with lectin (**Ai**) with segmentation of the arterial tree and organs overlayed (**Aii**). Removal of raw signal allows for the branching pattern of the arterial system supplying the bladder to be visualized (**Aiii**). The right side of the bladder has two VAs branching from the UA, giving rise to a DVA and DDA (**Aiv**). The left side of the bladder is supplied by one VA branching from the UA; this subsequently branches near the bladder, providing a dorsal branch supplying the DDA and a ventral branch supplying the SVLA and DVA (**Av**). **B**, Male pelvis labelled with lectin (**Bi**) with segmentation of the arterial tree and organs overlayed (**Bii**). Due to the positioning of glandular tissue leading to difficulties in imaging, the left internal iliac was unable to be identified in this image, leading to a gap in the segmentation. Removal of raw signal allows for the branching pattern of the arterial system supplying the bladder to be visualised (**Biii**). The right side of the bladder has one VA, which bifurcates forming a ventral branch supplying the DVA and a dorsal branch which subsequently bifurcates ventrally and dorsally supplying the SVLA and DDA, respectively (**Biv**). The left side of the bladder has one VA, which bifurcates into a ventral branch suppling the DVA and SVLA, and a dorsal branch suppling the DDA (**Bv**). Scale bars: 1000 µm.

Following the trajectory of the VAs, we were able to clearly identify the origin of the SVLA, DDA and DVA, however the smaller size of the DMA made it difficult to consistently visualize in these larger samples. The precise site of origin on the VA the urinary bladder arteries SVLA, DDA and DVA varied between animals. From analyses of vascular anatomy in both sides of adult mice, on the sides where there was one VA (n = 19), 5 different branching patterns were observed (Fig. 6C), two of which were more common (Fig. 6Ci**, 6Cii**). In one of the two most common patterns (n = 6 sides), the DDA originated most proximally to the urogenital artery and a bifurcation of the SVLA and DVA occurred most distally, i.e., closer to the bladder (Fig. 4Av, Fig. 4Bv, Fig. 6Ci). The other common pattern in mice with one VA had the more proximal branch of the vesicular artery branch near the bladder, penetrate the surface to then give rise to the DVA and the DDA, while the VA would continue to project towards the bladder where it would form the SVLA (n = 7 sides) (Fig. 6Cii). The final pattern with one VA and an SVLA was infrequent (n = 1); here the most proximal branch of the VA supplied only the DVA, while the SVLA and DDA originated from the bifurcation of the VA at the bladder surface (Fig. 4Biv, Fig. 6Ciii). In mice having two VAs supplying a side of the bladder, most commonly the VA more proximal to the internal iliac projected to the ventral side (supplying the SVLA and DVA) and the more distal VA projected to the dorsal supply (supplying the DDA) (n = 10 sides) (**Fig 6Civ**).

The most common pattern without an SVLA present (n = 7 sides) comprised two VAs branching from the urogenital artery, the more proximal of which projected to the ventral side of the bladder supplying the DVA, while the second VA projected to the dorsal side supplying the DDA (Fig. 4Aiv, Fig. 6Cvii).

The larger size of samples containing additional pelvic structures impaired more detailed analysis of the vasculature within the bladder wall. Therefore, to provide the optimal technical conditions for immunolabeling and microscopy of vascular structures on the surface or within the urinary bladder, most of the surrounding tissues were removed from the sample at the time of initial dissection or prior to processing of fixed samples. In these samples, the VAs and origins of the major arterial vessels from the VAs were not always present.

### Venous network and capillaries of the adult urinary bladder

Visualization of the venous supply was performed in both cleared intact bladders and flat mount preparations, identifying venous structures by their immunoreactivity for endothelial proteins (PECAM1/PODXL) but absence of lectin labelling. Each of the four main arterial supplies were examined for close association with veins. The SVLA (Fig. 5A) and DMA (Fig. 5B) were not closely associated with veins. PECAM1/PODXL immunolabeling was present in vessels near the SVLA and DMA but this was weak and patchy, the associated vessels subsequently identified as lymphatic vessels (Lyve-1). In contrast, the two deep arteries, the DVA and DDA, were closely associated with veins (Fig. 5C, D). A pair of interconnected veins running parallel along either side of the DVA and DDA periodically joined and separated. At branch points of the DVA and DDA, the associated veins also branched, then forming capillary beds. These primary relationships between the arterial and venous systems are represented in Fig. 6. These features were seen in both sexes.

**Fig. 5.**
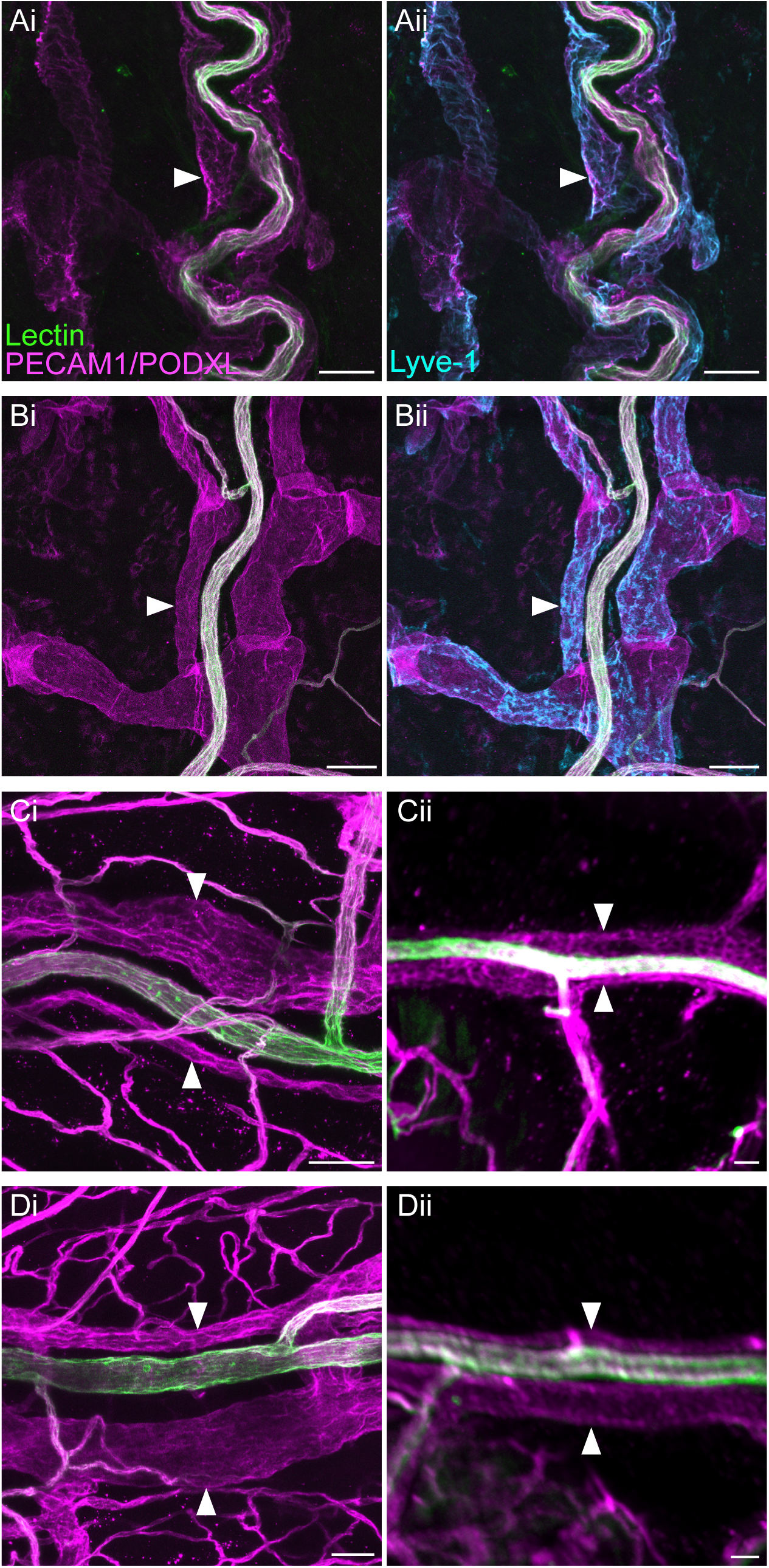
Intramural venous supply of the adult mouse urinary bladder. Confocal and light sheet microscopy of bladders dissected from female (**A, Ci, Di**) and male (**B, Cii, Dii**) mice and fixed as opened flat sheets (confocal microscopy; **A, B, Ci, Di**) or as intact organs (light sheet microscopy; **Cii, Dii)**. Intracardiac injection of fluorophore-conjugated tomato lectin preceded fixation and labelled the arterial system. The potential presence of closely associated veins was examined for each of the four main arterial supplies. Colocalization of lectin (green) and PECAM1/PODXL (magenta) is indicated as white. **A**, Superficial ventrolateral artery, accompanied on both sides by a vessel visualized by weak patchy immunoreactivity for PECAM1/PODXL (**Ai**). These are not veins but lymphatic vessels, indicated by Lyve-1 (**Aii**, arrow). **B**, Dorsal midline artery is also devoid of closely associated veins, but lymphatic vessels are nearby (Lyve-1, arrow). **C**, Deep ventral artery. Arrows indicate a vein each side of the artery, visualized using confocal (**Ci**) and light sheet (**Cii**) microscopy (20 µm optical slice). **D**, Deep dorsal artery. Arrows indicate a vein each side of the artery, visualized using confocal (**Di**) and light sheet (**Dii**) microscopy (20 µm optical slice). Scale bars: 50 µm.

**Fig. 6.**
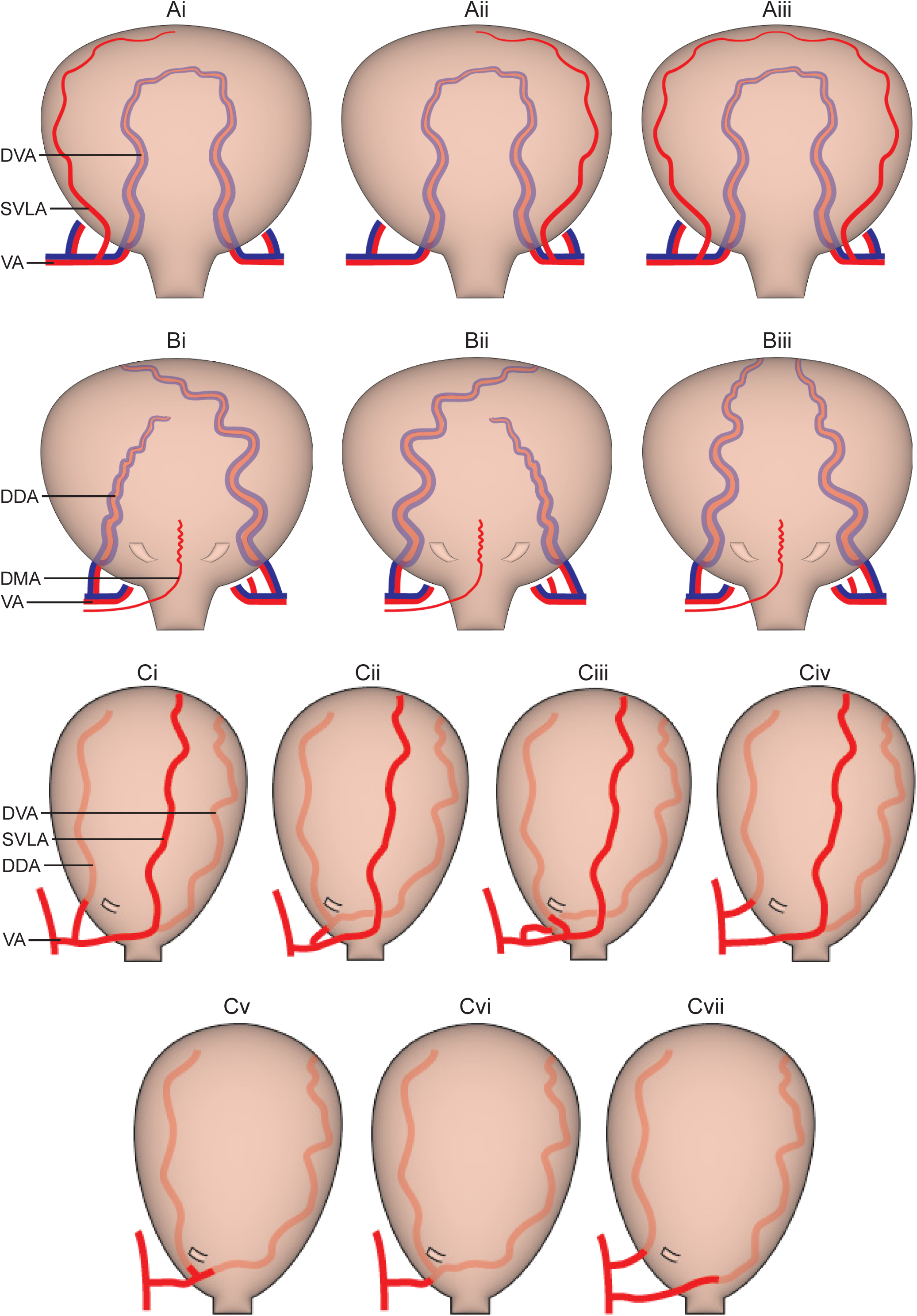
Schematic representation of the primary arterial and venous supplies of the adult mouse urinary bladder. DDA, deep dorsal artery; DMA, deep midline artery; DVA, deep ventral artery; SVLA, superficial ventrolateral artery; VA, vesical artery. Red, arteries; Blue, veins. **A**, Ventral side of the bladder, showing the variations in the superior ventrolateral artery (SVLA) that can be unilateral (either side; **Ai, Aii**) or paired (**Aiii**). **B**, Dorsal side of the bladder, showing variations in the deep dorsal artery (DDA) that can be unilateral (either side; **Bi, Bii**) or paired (**Biii**). The entry points of the ureters are shown. **C**, Lateral view of the bladder with the ventral side to the right, showing variations in the number of vesical arteries (VA; one VA **Ci, Cii, Ciii, Cv and Cvi;** two VA **Civ and Cvii**) and the branch points of the SVLA, DVA and DDA. These extramural vascular networks were assessed in both sides of 18 mice, including both sexes, where the prevalence of each structural class was: **Ci** (6), **Cii** (7), **Ciii** (1), **Civ** (10), **Cv** (2), **Cvi** (3), **Cvii** (7).

Lectin labelling of the capillary network was of variable intensity across samples so was not used for analysis of this network. To label the entire vascular network, the combination of PECAM1 and PODXL antibodies strongly labelled capillaries in intact cleared bladders and opened flat mounts. In the bladder we observed two distinct capillary beds, one within the muscularis and the other in the lamina propria (Fig. 7A). The capillary bed was much sparser in the muscularis (Fig. 7Aii) than the lamina propria (**Fig, 7Aiii**). These capillary beds were supplied by vessels perpendicular to the surface (Fig. 7Ai) which were either direct branches from the major arteries or branches from smaller arterioles. Some of the small arterioles supplying the lamina propria capillary bed demonstrated a constriction at their branch points (Fig. 7B); these may correlate with the arterial sphincters described previously (7). Using cleared samples of the entire LUT, we were also able to identify that both the muscularis and lamina propria capillary networks of the urinary bladder were continuous with those of the adjacent urethra and ureter (**Supplementary Video S5** https://doi.org/10.26188/28844171.v1 ).

**Fig. 7.**
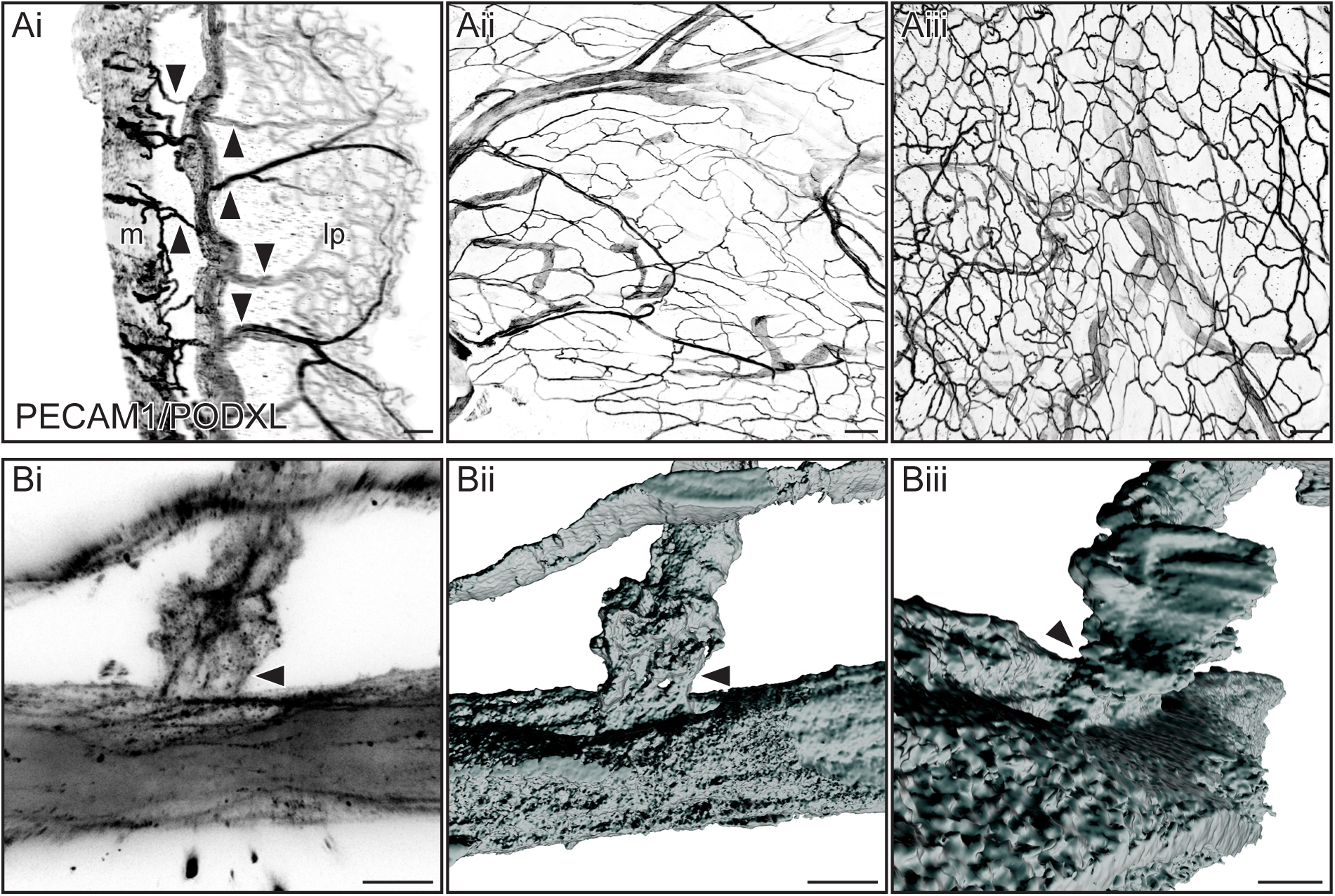
Capillary network of the adult mouse urinary bladder. Confocal and light sheet microscopy of bladders dissected from female mice and fixed as intact organs (light sheet microscopy; **Ai**) or opened flat sheets (confocal microscopy; **Aii, Aiii, B**). PECAM1/PODXL immunolabeling was used to visualize capillary networks. m, muscularis; lp, lamina propria **A**, Capillary networks in the muscularis and lamina propria. **Ai**, optical slice (200 µm) from light sheet image, demonstrating the muscularis and lamina propria capillary beds originating from the deep ventral artery and its associated veins projecting near the muscularis-lamina propria boundary. Arrows indicate perpendicular vessels arising from the supplying artery and veins. The different density of the capillary beds in each tissue is shown more clearly in flat mounts (**Aii**, muscularis; **Aiii**, lamina propria). **B**, An example of an arteriole that supplies capillaries in the lamina propria exhibiting a constriction at a branch point, potentially indicating an arterial sphincter (arrows), To visualize this feature more clearly, segmentation using Imaris Surfaces was performed, shown here in two different orientations. Scale bars: A 100 µm, B 10 µm.

### Blood vascular supply of the immature urinary bladder

Similar studies were performed in immature mice (P7-P10) to determine which features of the vascular structure were already established at this stage. In these smaller animals, perfusion of lectin after intracardiac injection was typically compromised due to a rapid weakening or cessation of the heartbeat. This problem was partially overcome by reducing the lectin volume from 100 µl to 20-50 µl, however in most immature animals the lectin signal remained weaker and less complete than in adult mice. Therefore, immunolabeling with PECAM1/PODXL was included in all samples; this was critical for visualization and validation of all structures.

The four major arteries identified in adult mice (SVLA, DDA, DVA, DMA) were established by P7. Their primary features (e.g., location within the bladder and region supplied) appeared comparable to adults. Only one difference was identified, relating to the number of SVLAs. In the male pup bladder, 2 SVLAs were present in all 15 preparations (Fig. 8A), whereas in adult males, only 3 of 14 preparations had bilateral SVLAs. This increased prevalence of 2 SVLAs in pups was also identified in females, where 2 SVLAs were present in 13/16 female pups but only 7/18 female adults. In the 3 female pups with only one SVLA, this vessel was always on the left side.

**Fig 8:**
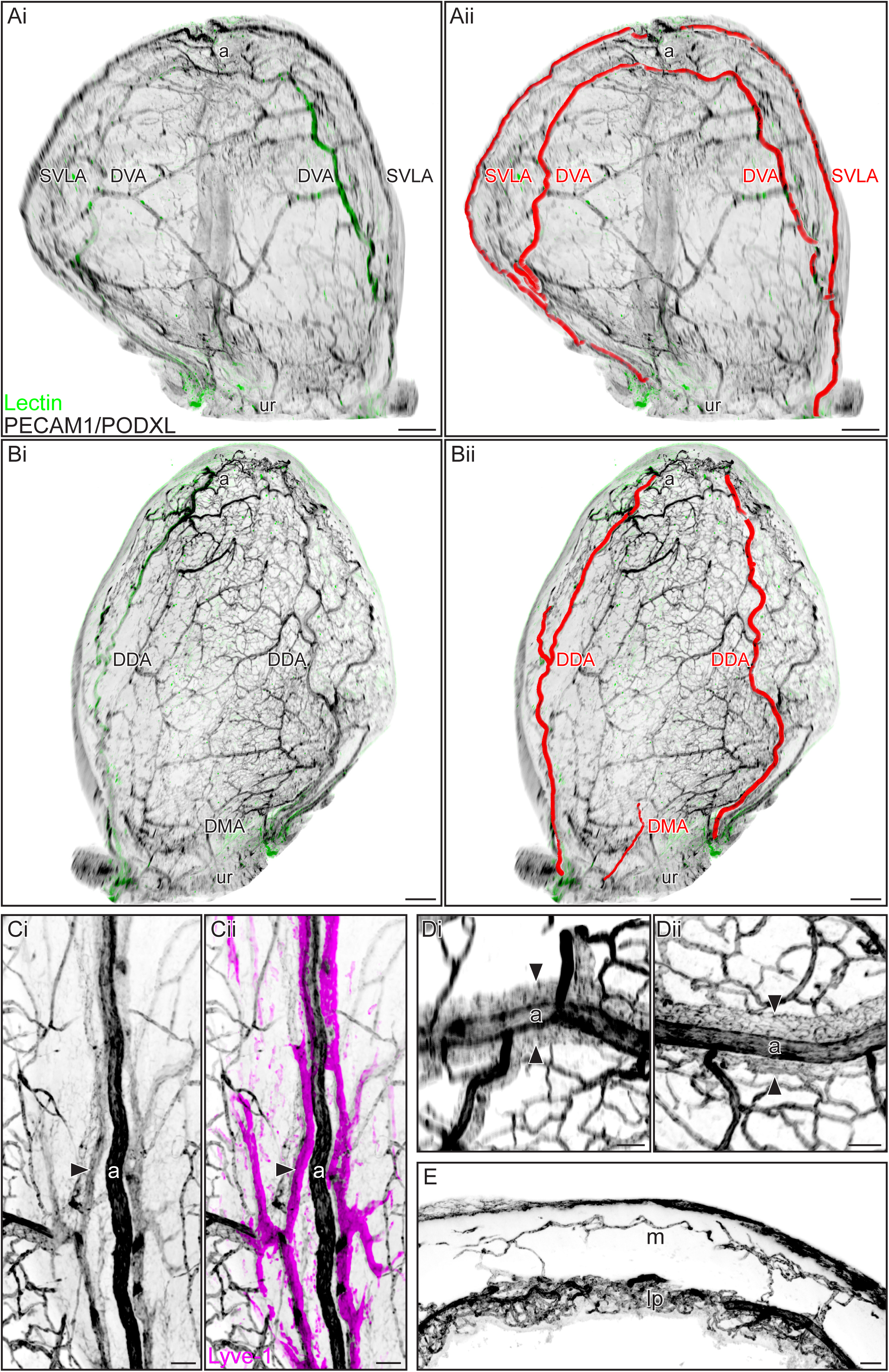
Vascular supply of the immature urinary bladder. Light sheet microscopy of whole intact bladders from male (**A, B, D and E**) and female (**C**) mouse pups (postnatal days P7 A-B; P8 C-E). Lectin labelling was generally weaker and less complete in these immature animals, so the digital colors have been chosen to emphasise the PECAM1/PODXL immunolabeling that demonstrates the complete vascular network. All segmentations were performed with Imaris. Monochrome images have been inverted to enable better visualization of vascular structures. a, apex of bladder; ur, urethral orifice. **A**, Ventral view of the bladder, showing the superficial ventrolateral arteries (SVLAs) and deep ventral arteries (DVAs). Original image (**Ai**) shows the SVLAs, DVAs and many other smaller vessels from the deeper vascular network, with **Aii** showing the segmented SVLAs and DVAs. **B**, Dorsal view of the bladder, showing the deep dorsal arteries (DDAs and DMA). Original image (**Bi**) shows the DDAs and many other smaller vessels from the deeper vascular network, with **Bii** showing the segmented DDAs and DMA. **C**, Superficial ventrolateral artery (a) accompanied on both side by vessels with weakly positive PECAM1/PODXL labelling (**Ci**). These were defined as lymphatic vessels, as indicated by Lyve-1 labelling (**Cii**, 100 µm optical slice). **D**, Deep ventral (Di) and dorsal (Dii) arteries (a) were accompanied on both sides by veins, indicated by arrows in two samples (100 µm optical slices). **E**, Capillary beds of the two layers of the bladder, highlighting the higher density of the capillary bed within the lamina propria (lp) compared to the muscularis (m). Visualised using a 200 µm optical slice. Scale bars: A-B 300 µm, C-E 50 µm.

The remaining major arteries (DDA, DVA, DMA) were also similar between pups and adults. The DVAs in pups had the characteristic hair-pin structure (Fig. 8A). As observed in some adult mice, in pups it was common that one DDA supplied a greater area of the dorsal side of the bladder (Fig. 8B). This regional dominance was identified in 8/16 female pups (3 showed right and 5 left dominance). In male pups, 4 samples displayed regional dominance (2 right and 2 left). Even when lectin labelling was optimal, we could not identify the DMA in all preparations. Of the 16 females, the DMA was present in 15; of these, the DMA originated from the right sidein 8 samples and the left in 7 samples. Of the 15 males, 14 had labelling of sufficient quality in this region; here, the DMA was present in 12/14 pups and of these, the DMA branched from the right side in 8 samples and the left in 4 samples.

The other vascular structures associated with these major arteries in adults were established by P7. There was no vein closely associated with the SVLA, but instead a lymphatic vessel was closely aligned with each side (Fig. 8C). Also comparable to adults, the DDA and DVA were each associated with a pair of veins; these veins travelled close to the relevant artery for most of their length (Fig. 8D). Two independent capillary beds belonging to the muscularis and lamina propria were also evident (Fig. 8E), and as seen in the adults, the capillary bed was denser in the lamina propria than the muscularis.

## Discussion

In this study we have identified stereotypical structural features of the arterial network associated with the urinary bladder in adult male and female mice. To our knowledge, these features have not yet been defined in the rodent urinary bladder or in other species. This new understanding of stereotypical patterning and specific sites of origin for the vascular supply of different bladder tissues and regions will inform experimental design, analysis and modeling of bladder vascular function and pathophysiology.

Current understanding of vascular structures within the urinary bladder is strongly based on structural studies using resin corrosion casting and scanning electron microscopy (7–11, 31) and physiological studies on specific components of the microvasculature (5, 6, 13, 32, 33). The resin casting approach has been applied to the adult urinary bladder in several species (dog, rabbit, mouse, human) and revealed distinct features of capillary networks and specializations such as coiled trajectories, anastomoses and valves (in veins). Our primary approach of tissue clearing with light sheet microscopy to view the entire bladder has enabled analysis of this vasculature across scale – from the large vessels that comprise the extramural sources of bladder arteries to the capillaries and other microscopic structural features. Supplementing this approach with opened, flat whole mounts has enabled confirmation of structures visible with light sheet microscopy and higher resolution visualization.

This multi-scale approach has enabled us to identify a robust, stereotypical anatomy of the bladder arterial system and its relationship to specific regions and tissues, in both sexes and across postnatal developmental stages, in addition to its relationship to the venous and lymphatic systems. Our analyses were facilitated by an intravital labeling protocol that selectively visualized the arterial system. This increased the efficiency of identifying and tracing the primary arterial networks without the visual distractions of the venous system. We identified four major arteries associated with the bladder wall (superficial ventrolateral artery [SVLA], deep dorsal artery [DDA], deep ventral artery [DVA] and dorsal midline artery [DMA]), each of which had specific trajectories, regional and tissue supplies. We have provided new, anatomically descriptive names for these vessels as we found no previous reports of all four vessels in the literature. From the images and descriptions provided, we consider it likely that the SVLA is the same vessel identified in the rabbit bladder as the serpentine vesicular artery (9); in this same study, another vessel was described as ‘’a large straight branch supplying the base of the bladder”, which may be the DMA identified by the current study.

It is not known if each of these arterial supplies are impacted differently by experimental perturbations or participate differently in adaptive or regenerative responses to challenge. Single cell transcriptome analysis of mouse and human organs has revealed organ-specific features of endothelial cells and pericytes within the vasculature (34–36). It would be interesting to investigate if the four major arterial systems or their associated capillaries and veins have distinct molecular and regulatory features. We further propose investigation of whether stereotypical intramural arterial systems exist within the human bladder; if so, this knowledge would benefit many aspects of urological, neuro-urological, surgical and bladder cancer studies.

Microscopy of cleared intact pelvis samples allowed us to visualize the origin of the four major arterial vessels of the bladder. These large, complex samples also contain the vascular supply of many other pelvic organs and tissues that were not examined further here, although an additional study on urethral vasculature is underway. In the context of bladder vasculature, these samples showed several anatomical variations in branching location and number, but all originated from one or two vesical arteries, in turn originating from the urogenital artery and anterior branch of the internal iliac (noting that the source of the DMA was generally more difficult to identify and may be more variable). This aligns with extramural vascular patterning reported by a previous study in the wood mouse (37) although this study also reported variations in the origin of the urogenital and internal iliac arteries. A study in “black laboratory mice” reported similar origins and branching patterns but did not identify animals with two vesicular arteries (7).

The laminar location of the primary trajectory of each artery was either superficial, i.e. associated with the serosa (SVLA, DMA) or deeper within the bladder wall, specifically at the boundary of the muscularis and lamina propria (DDA, DVA). The SVLA and DMA branch to supply capillaries to adjacent tissues, however whereas the SVLA contributed to the capillary beds of the muscularis and lamina propria, the DMA only appeared to supply the muscularis. However, it should be noted that the more challenging imaging of the DMA may have underestimated the extent of its derived vascular network. We observed a highly tortuous termination of the SVLA at the apex, where it provided a dense capillary network. This suggests a unique type of vascular function at this location, as also proposed by Hossler and colleagues (9) who visualized this complexity at the apex of the rabbit bladder and suggested that this very dense complex of vessels at the apex may accommodate bladder distension. We are unaware of studies specifically investigating the function of these vessels near the apex or, for other bladder regions, whether each of the four main arterial networks contribute specifically to overall vascular perfusion (or if they respond similarly to challenge). The DDA and DVA both supplied capillary beds in the muscularis and lamina propria.

Although our primary focus was the arterial system, we were also able to visualize the complete venous and capillary networks. As reported previously, the capillary beds in the lamina propria and muscularis were quite distinct, with the former comprising a much denser network, indicating higher metabolic requirements (9, 11, 38–40). We were also able to view the continuity in capillary networks between the bladder and the ureter and urethra, as previously reported in mice (7). These junctions or transitions between organs may be important sites of change during developmental or experimental perturbations but are only visible in these intact preparations.

Each of these vessels has a distinct relationship to the venous or lymphatic systems, with the two deep arteries (DDA, DVA) having a pair of closely associated veins. The trajectory of these deep arteries and associated veins across the bladder wall occurs at the boundary of the muscularis and lamina propria, with branches emanating on both sides to supply capillary beds of both tissues. In contrast, the more superficially located SVLA and DMA were not closely associated with veins but instead the SVLA was located closely between a pair of large lymphatic vessels. The close apposition between the now identified and named SVLA and lymphatic vessels is clearly visible in a published image of whole mount mouse bladder (41), however to our knowledge the functional relevance of this relationship has not been explored. A study specifically investigating this relationship and the complex lymphatic network of the LUT is underway. Vessels closely aligned with the serpentine vesicular artery in rabbits (potentially a homologue of the SVLA in mice) were likely to be veins rather than lymphatic vessels as they filled with resin during the vascular casting process (9).

The major features of the vascular system identified in our study are established by the early postnatal period (P7-P10). At this time the neural pathways underpinning voiding and continence are not yet fully mature, with volitional voiding only being established after weaning (1). However, we did identify a difference in laterality of vascular networks between immature mice (P7-10) and adults. This was most evident for the SVLA that was unilateral in the majority of adult mice but was more commonly bilateral in pups; at both ages, in most animals where the SVLA was unilateral, the SVLA was present on the left side. We infer that the second SVLA regresses between P10 and adulthood but have not investigated this timing further. The termination of the SVLA at the apex raises the possibility of an earlier connection between the SVLA and the umbilical artery, which to our knowledge has not yet been examined. The mechanisms driving this laterality and proposed tissue regression should also be explored.

We did not observe sex differences in the features of the bladder documented here, however our study did not quantify specific structural properties of the vasculature, such as branching patterns, vessel diameter or capillary density in each region. We are currently utilizing the 3D datasets from this study to build computational pipelines for this purpose. It is possible these will reveal regional, tissue and sex differences. We are also performing a detailed analysis of the vascular system in the urethra, using a similar approach to the current study.

In experimental studies on the bladder, it has been difficult to assess or communicate regional changes in bladder function with the existing range of anatomical terms for this organ (e.g., bladder apex, dome, base, neck) that do not all have distinct anatomical boundaries and are inconsistently applied. This has restricted progress on integrating data across experimental studies or interpreting new results in the context of past work. The same limitations apply to clinical studies on the urinary bladder. Studies on other organ systems have benefited from the development of a common coordinate framework (CCF) (42), however to our knowledge a CCF has not yet been established for the lower urinary tract of any species. The underlying vascular network structure within an organ can provide a robust foundation for building such a framework (43), especially where there are few other anatomical landmarks to indicate specific regions. We propose that the four main arterial supplies identified in the current study could form an important component of a new bladder CCF. We further propose that clinical anatomical studies investigate whether stereotypical intra-mural arterial systems exist within the human bladder, knowledge that would benefit many aspects of urological, neuro-urological, surgical and bladder cancer studies.

## Conclusion

This study has identified stereotypical arterial and venous networks associated with the mouse urinary bladder and showed that the primary features of this network are established by the early postnatal period. This new anatomical knowledge will inform studies on physiological and pathophysiological changes in the urinary bladder vascular network and facilitate more refined experimental perturbation, analysis and interpretation of vascular function/dysfunction in mouse models.

## Acknowledgements

We acknowledge the Biological Optical Microscopy Platform (BOMP), University of Melbourne (RRID:SCR_018888), for providing training and access to the light sheet microscope and confocal microscopes. Dr Ilvana Ziko and Mr Dain Maxwell provided technical support for the study, including assistance with animal procedures and tissue processing. Research reported in this publication was supported by the National Institutes of Health under Award Number U01DK131384. The content is solely the responsibility of the authors and does not necessarily represent the official views of the National Institutes of Health.

## Author contributions

JK, PO and LB Conceived and designed research; LB and MD Performed experiments, interpreted results of experiments and analyzed data; LB prepared figures; LB and JK drafted the manuscript; All authors edited, revised and approved the final version of the manuscript.

## Supplementary videos S1-S5

**Supplementary video S1: Major intramural arteries of the adult mouse urinary bladder**

https://doi.org/10.26188/28844174.v1

Cleared whole lower urinary tract from an adult female mouse, perfused intracardially with a DY649 conjugate of Tomato lectin and immunostained for vascular markers PECAM1 and PODXL and viewed with light sheet microscopy (Ultramicroscope II, Miltenyi Biotec, Germay). Segmentations (Imaris) of the superficial ventrolateral artery (SVLA), deep ventral arteries (DVA) and deep dorsal arteries (DDA). This demonstrates the stereotypical branching and location of these major arteries.

**Supplementary video S2: Dorsal Midline Artery of the adult mouse urinary bladder**

https://doi.org/10.26188/28844177.v1

Cleared whole lower urinary tract from an adult male mouse, perfused intracardially with a DY649 conjugate of Tomato lectin and immunostained for vascular markers PECAM1 and PODXL and viewed with light sheet microscopy (Ultramicroscope Blaze, Miltenyi Biotec, Germany). Image shows the dorsal side of the bladder, including the bladder neck. Some of the surrounding tissues are also seen (prostate, seminal vesicles, vas deferens). Segmentation (Imaris) and red arrows highlight the dorsal midline artery (DMA).

**Supplementary video S3: Extramural vascular supply to the female mouse urinary bladder**

https://doi.org/10.26188/29083577.v1

Cleared lower abdomen and pelvis of adult female mouse perfused, intracardially with a DY649 conjugate of Tomato lectin and viewed with light sheet microscopy (Ultramicroscope Blaze, Miltenyi Biotec, Germany). Segmentations of vessels (Syglass) and organs (Imaris) demonstrating the origins of the vasculature supply of the urinary bladder in the context of the entire pelvis. Red = vessels of the external supply (abdominal aorta, common iliac, internal iliac, urogenital artery); orange = vesicular arteries; yellow = intramural vessels of the bladder, i.e., deep dorsal artery (DDA), deep ventral artery (DVA) and superficial ventrolateral artery (SVLA).

**Supplementary video S4: Extramural vascular supply to the male mouse urinary bladder**

https://doi.org/10.26188/29062364.v1

Cleared lower abdomen and pelvis of adult male mouse perfused, intracardially with a DY649 conjugate of Tomato lectin and viewed with light sheet microscopy (Ultramicroscope Blaze, Miltenyi Biotec, Germany). Segmentations of vessels (Syglass) and organs (Imaris) demonstrating the origins of the vasculature supply of the urinary bladder in the context of the entire pelvis. In this visualization the left internal iliac is not shown. Red = vessels of the external supply (abdominal aorta, common iliac, internal iliac, urogenital artery); orange = vesicular arteries; yellow = intramural vessels of the bladder, i.e., deep dorsal artery (DDA), deep ventral artery (DVA) and superficial ventrolateral artery (SVLA).

**Supplementary video S5: Relationships between the vascular networks of the mouse urinary bladder, ureter and urethra**

https://doi.org/10.26188/28844171.v1

Cleared whole lower urinary tract from adult female mouse immunostained for vascular markers PECAM1 and PODXL and viewed with light sheet microscopy (Ultramicroscope II, Miltenyi Biotec, Germany). Image shows the bladder neck with a short length of ureter and proximal urethra attached. Autofluorescence was used to visualize the organs and shows that vascular networks were continuous between them (visualized using Imaris). Red arrows indicate a blood vessel that projects from the urethra via the bladder neck into the bladder wall.

